# Trunk Control during Gait: Walking with Wide and Narrow Step Widths Present Distinct Challenges

**DOI:** 10.1101/2020.08.30.274423

**Authors:** Hai-Jung Steffi Shih, James Gordon, Kornelia Kulig

## Abstract

The active control of the trunk plays an important role in frontal plane gait stability. We characterized trunk control in response to different step widths using a novel feedback system and examined the different effects of wide and narrow step widths as they each present unique task demands. Twenty healthy young adults walked on a treadmill at 1.25 m/s at five prescribed step widths: 0.33, 1.67, 1, 1.33, 1.67 times preferred step width. Motion capture was used to record trunk kinematics, and surface electromyography was used to record longissimus muscle activation bilaterally. Vector coding was used to analyze coordination between pelvis and thorax segments of the trunk. Results showed that while center of mass only varied across step width in the mediolateral direction, trunk kinematics in all three planes were affected by changes in step width. Angular excursions of the trunk segments increased only with wider widths in the transverse plane. Thorax-pelvis kinematic coordination was affected more by wider widths in transverse plane and by narrower widths in the frontal plane. Peak longissimus activation and bilateral co-activation increased as step widths became narrower. As a control task, walking with varied step widths is not simply a continuum of adjustments from narrow to wide. Rather, narrowing step width and widening step width from the preferred width represent distinct control challenges that are managed in different ways. This study provides foundation for future investigations on the trunk during gait in different populations.

## Introduction

Maintaining stability is one of the primary goals in human locomotion. In contrast with sagittal plane stability, which is largely achieved through passive mechanisms, frontal plane stability during gait requires active control (Bauby and Kuo, 2000; Kuo and Donelan, 2010). Frontal plane stability during walking is achieved through a combination of different strategies, including adjusting mediolateral motion of the body’s center of mass (CoM) (Arvin et al., 2016a), manipulating mediolateral foot placement (Bruijn and van Dieën, 2018) or external rotation of the foot (Rebula et al., 2017). The trunk accounts for 48% of the body mass and is the largest contributor to the CoM (MacKinnon and Winter, 1993; Prince et al., 1994). Therefore, how the trunk is controlled will directly influence the CoM, and in some cases adjustment of the CoM motion is preferred over modifying foot placement (Best et al., 2019). The trunk should not be considered a rigid body, since structurally it involves multiple linked segments and degrees of freedom that needs to be fine-tuned. When these degrees of freedom within the trunk are constrained with an orthosis, an impact on CoM excursion and step width is observed (Arvin et al., 2016b).

Although the relationship between CoM and lateral foot placement has been extensively studied (Arvin et al., 2016a; Bruijn and van Dieën, 2018; Hurt et al., 2010; McAndrew Young and Dingwell, 2012; Perry and Srinivasan, 2017; Stimpson et al., 2018; Wang and Srinivasan, 2014), the within-trunk control in response to step width has not been well-investigated. Furthermore, the use of strings or tape to prescribe step width in previous studies required participants to look down, affecting their trunk motion (Arvin et al., 2016a; Perry and Srinivasan, 2017). We therefore designed a study using visual feedback at eye-level to investigate trunk control at different step widths. This study will provide insight into walking mechanics and a basis for understanding and training trunk motion in different populations, such as persons with spinal pathology, balance issues, or older adults.

During gait, the mass of the trunk is controlled in the frontal plane mainly by the spinal musculature and the hip abductors (MacKinnon and Winter, 1993). Studies have primarily focused on the contribution of hip abductors to modifying foot placement (Rankin et al., 2014) and stability during stance phase (Kubinski et al., 2015). However, the role of paraspinal muscle activation has not received much research attention. A study on typical walking demonstrated that paraspinal muscles at the lumbar region were the most highly activated among the C7 to L4 paraspinal muscles (Prince et al., 1994), but how the muscles’ activation patterns are modulated for different step widths is still unknown.

Walking with wider and narrower step widths places unique demands on the motor system. The preferred step width is likely to be selected to minimize energy cost without compromising stability (Donelan et al., 2001). Moving away from the preferred step width affects the inverted pendulum-like motion of the CoM and influences energy demands (Kuo et al., 2005). A previous study demonstrated that walking with wider step widths requires greater mechanical work to redirect the CoM, while walking with narrower step widths increases work associated with swinging the leg laterally to avoid the stance limb (Donelan et al., 2001). Walking with a narrower width also presents greater challenges to stability and increases demand on active postural control as the base of support decreases (Donelan et al., 2004; MacKinnon and Winter, 1993; Perry and Srinivasan, 2017). The current study will investigate whether trunk control varies continuously across step widths or presents as separate motor patterns in wide and narrow widths. If the latter is true, walking with wide and narrow widths can be viewed as two distinct tasks.

The purpose of this study was to characterize changes in trunk control during different prescribed step widths, and to compare the effects of wide and narrow step widths on trunk kinematics and muscle activation. We hypothesized that narrower widths will present greater stability demands and result in an increase in muscle co-activation, while wider widths will present greater mechanical demands and influence trunk kinematics.

## Methods

### Participants

Twenty healthy young adults participated in this study (14 females, 6 males; 26.25 ± 3.31 years; 165.54 ± 9.93 cm; 61.39 ± 12.71 kg; BMI = 22.21 ± 2.84 kg/m2). Participants were included if they were between 18 to 45 years old and excluded if they had a history of lower extremity or spine pathology or surgery. Participants gave written informed consent that was approved by the university’s institutional review board.

### Instrumentation

Participants were instrumented with a lower extremity marker set and additional markers on bilateral acromion, sternal notch, and T1. Kinematic data were recorded by a 11-camera Qualisys motion capture system (Qualisys, Gothenburg, Sweden) at 125 Hz. Surface electrodes were placed on bilateral longissimus (2 finger widths lateral from the spinous process of L3). Electromyography (EMG) data were collected using Noraxon wireless EMG system (Noraxon U.S.A, Scottsdale, AZ, USA) at 1500 Hz. A portable treadmill (ICON Health & Fitness, Logan, UT, USA) was used for the walking trials.

### Experimental procedures

Participants completed a medical history form and the International Physical Activity Questionnaire (Craig et al., 2003). Leg length from the greater trochanter to the ground was measured. A standing calibration trial for the kinematic data was collected. Participants were given up to 3 minutes to familiarize with the treadmill, after which a 30 second treadmill walking trial at 1.25 m/s was collected to determine preferred step width (PSW).

Participants then walked with 5 different step widths using real-time visual feedback projected on the wall in front of the treadmill (Fig 1). Step width was calculated using marker data that was streamed real-time and displayed as visual feedback through custom code written in MATLAB (MathWorks, Natick, MA, USA). The midpoint between the heel and 2nd toe marker positions at foot flat was calculated to indicate foot position, then the medial-lateral distance between subsequent foot falls were calculated as step width. This accounted for external rotation angle in addition to foot placement, since foot angle has an effect on base of support and plays a role in maintaining stability during walking (Rebula et al., 2017). Five different step widths were prescribed, including 0.33, 0.67, 1, 1.33, and 1.67 × PSW. One 30-second familiarization trial at 1PSW was given, followed by four 30-second practice trials, one trial for each prescribed width other than 1PSW. Participants then completed one 30-second trial for each step width in a randomized order.

**Figure 1.**
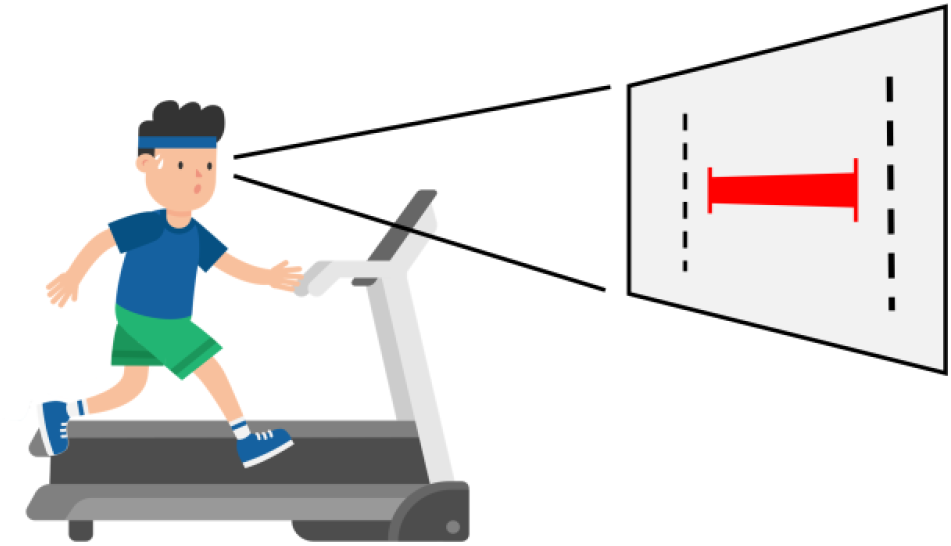
Experimental setup of treadmill walking with prescribed step widths. A visual feedback was projected on a wall in front of the treadmill, with a red horizontal bar representing participant’s actual step width and black vertical lines indicating the target width.

### Data Analyses

Kinematic data were low-pass filtered at 10 Hz with a dual-pass 4th order Butterworth filter. Gait events were identified based on the toe and heel marker position relative to the pelvis coordinate. Each gait cycle was time-normalized to 101 data points from right heel strike to the subsequent right heel strike. Each 30-second trial consists of approximately 25-35 strides, but only the last 20 strides were included in the analyses to allow participants to reach the prescribed step widths.

Step width was calculated as the mediolateral distance between foot positions of two successive foot falls. Constant step width error was calculated as the difference between the mean performed step width and the prescribed step width, and variable step width error was calculated as the standard deviation of the performed step width.

The thorax and pelvis angles were defined as the thorax/pelvis segment relative to the lab coordinate system, whereas the trunk angle was defined as the thorax relative to the pelvis (Fig 2A). Angular excursions were calculated as the peak-to-peak angles of the thorax, pelvis, and trunk during each gait cycle. Thorax-pelvis kinematic coordination was calculated using vector coding analysis (Needham et al., 2014). An angle-angle diagram was first constructed with the pelvis and thorax angles on each axes, then the coupling angle was determined as the angle of the vector between two adjacent data points in time relative to the right horizontal (Fig 2B). Coordination patterns were categorized as in-phase, anti-phase, thorax-only, and pelvic-only defined by coupling angles falling within each range illustrated in Fig 2C&D. Finally, the percentage of time in gait that participants spent in each pattern was calculated.

**Figure 2.**
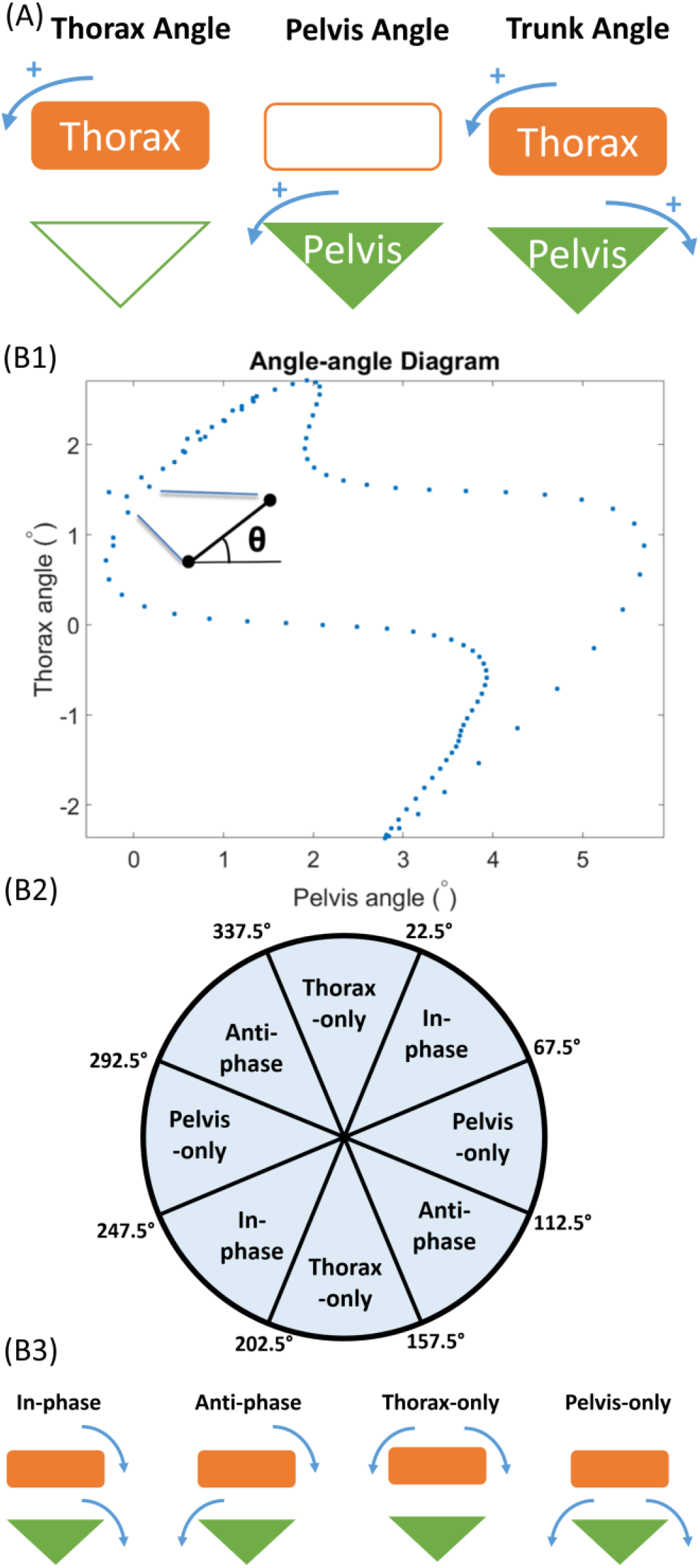
(A) Conceptual graphs illustrating our definition of thorax, pelvis, and trunk angles. Thorax angle and pelvis angles were defined as the thorax or pelvis segment angles relative to the lab’s coordinate system, while the trunk angle is the thorax relative to the pelvis segment angle. Showing frontal plane for ease of illustration, but these definitions also apply to the transverse and sagittal planes. (B1) Demonstration of vector coding analysis on an angle-angle diagram of a gait cycle in one representative participant. Coupling angle was defined as the vector angle of two consecutive points in time relative to the right horizontal. (B2) Cutoffs for binning of the coupling angles into four coordination patterns. (B3) Illustrations of the physical implication of the four coordination patterns. In-phase indicate that both segments are rotating towards the same direction at similar velocity; anti-phase indicate that the segments are rotating to the opposite direction at similar velocity; thorax-only and pelvis-only indicate that the thorax or pelvis segment is rotating significantly faster than the other segment, while the other segment may be hardly rotating.

EMG data were bandpass filtered between 30 Hz and 500 Hz with a dual-pass 4th order Butterworth filter to avoid heart rate contamination (Drake and Callaghan, 2006). The signal was then rectified and smoothed using a 100 ms moving window and normalized to the averaged peak activation during treadmill walking without step width prescription. Peak lumbar longissimus activation for each muscle was determined during the contralateral stance phase, where higher activation was found comparing bilateral stance phase. Bilateral longissimus coactivation was first determined at every sampling interval as the ratio of the less-activated side over the more-activated side at that sample point, then the summed ratio is divided by the number of samples across gait cycle. The following equation was used:

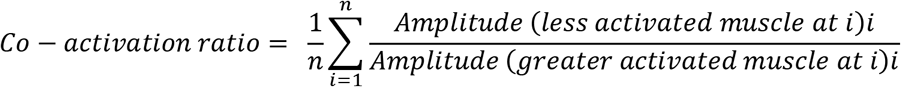

### Statistical Analyses

Data were plotted for visualization and examined for normality, homoscedasticity, and outliers. Descriptive statistics was performed on all variables. Statistic parametric mapping (SPM) was used to compare the time series data of trunk kinematics and detect phases in gait that were significantly different across step widths. In this study, the SPM F-statistic, a vectorfield equivalent of one-way repeated measures analysis of variance (ANOVA), was used. For more detailed information on SPM, see Pataky (Pataky, 2012, 2010). Constant and variable step width errors, thorax, pelvis, and trunk angular excursions, thorax-pelvis kinematic coordination patterns, lumbar longissimus EMG activity and bilateral co-activation were compared between step widths using traditional one-way repeated measures ANOVA. Tukey’s post-hoc tests were performed for multiple pairwise comparisons if the ANOVA F-test was significant. Analyses were done in MATLAB (Mathworks, Natick, MA) and R (R Core Team, 2018).

### Test-retest reliability

Six participants were tested again one week later to determine test-retest reliability. Test-retest intraclass correlation coefficients for one-way random effects, absolute agreement, and multiple measurements (ICC (1,k)) were calculated for all variables (McGraw and Wong, 1996). Additionally, standard error of the mean (SEM) was calculated as 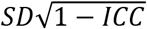. The α level was set at 0.05. Preferred step width, constant step width error, variable step width error, anti-phase and thorax-only coordination, peak longissimus activation, and bilateral longissimus co-activation had *good* test-retest reliability, and thorax, pelvis, and trunk angular excursion, inphase and pelvis-only coordination had *excellent* reliability (Table 1).

**Table 1.** Test-retest reliability and SEM for primary variables (ICC: intra-class correlation coefficient, SD: standard deviation, SEM: standard error of the measurement).

|  | ICC | SD | SEM |
| --- | --- | --- | --- |
| **Preferred Step Width (cm)** | 0.85 | 2.56 | 1.00 |
| **Constant Step Width Error (cm)** | 0.81 | 1.19 | 0.52 |
| **Variable Step Width Error (cm)** | 0.85 | 0.54 | 0.21 |
| **Thorax Angular Excursion (degree)** | 0.97 | 0.85 | 0.15 |
| **Pelvis Angular Excursion (degree)** | 0.93 | 2.21 | 0.57 |
| **Trunk Angular Excursion (degree)** | 0.94 | 2.52 | 0.61 |
| **In-phase Coordination (%)** | 0.93 | 8.05 | 2.13 |
| **Anti-phase Coordination (%)** | 0.78 | 3.73 | 1.75 |
| **Thorax-only Coordination (%)** | 0.85 | 3.99 | 1.57 |
| **Pelvis-only Coordination (%)** | 0.93 | 12.22 | 3.32 |
| **Peak Longissimus Activation (%)** | 0.73 | 5.38 | 2.77 |
| **Bilateral Longissimus Co-activation** | 0.76 | 0.03 | 0.02 |

## Results

### Task Performance

The average preferred step width (PSW) was 14.77 ± 2.92 cm, equivalent to 0.17 x leg length. Participants were able to match the different target step widths using the visual feedback. They made small but consistent positive errors that were dependent on step width, with the largest errors occurring at the narrowest step widths (Fig 3A). There was a significant difference in constant step width error between step widths (F(4,76) = 18.93, p<0.001) (Fig 3B). Post-hoc test revealed a significantly increased error at 0.33PSW compared to all of the other widths (p<0.003), and an increased error at 0.67PSW than 1.67PSW (p<0.001). There was also a significant difference in variable step width error between step widths (F(4,76) = 4.65, p=0.002), with significantly increased error at the 0.33PSW compared to 0.67, 1, and 1.33PSW (p<0.014).

**Figure 3.**
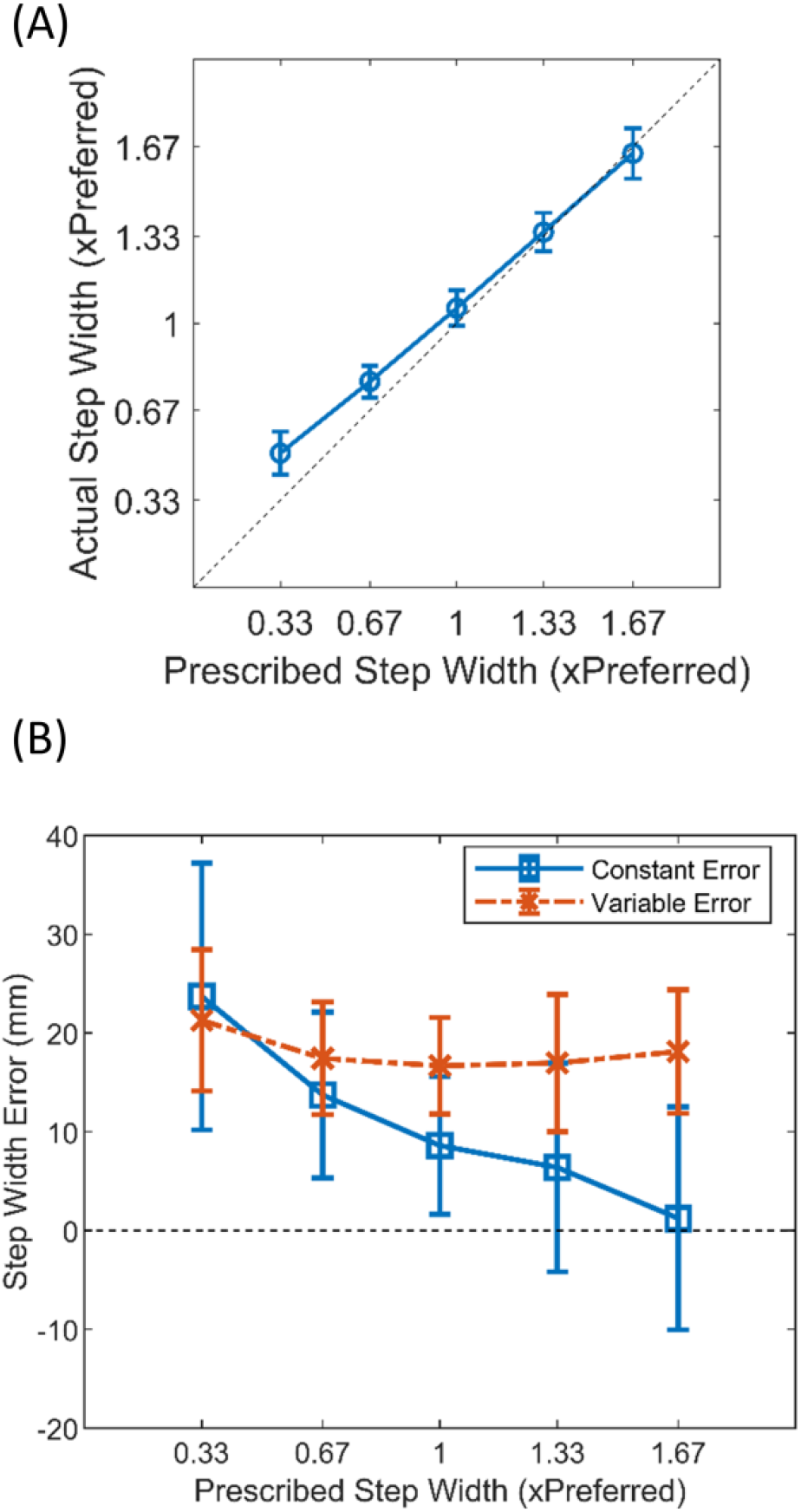
Step width task performance. (A) Square plot presenting the mean and standard deviation of actual step width performance relative to the prescribed step widths. The diagonal reference line indicates a one-to-one fit of the performance with the targets. (B) Mean and standard deviation of constant and variable step width error. Step width affected constant error more than variable error.

### Center of Mass and Thorax, Pelvis, and Trunk Kinematics

The amplitude of mediolateral center of mass (CoM) deviation scaled with step width, while the average anteroposterior and vertical CoM trajectories were hardly affected (Fig 4). Nevertheless, walking with different step widths resulted in different patterns of thorax, pelvis, and trunk angular motion in all three planes (Fig 5).

**Figure 4.**
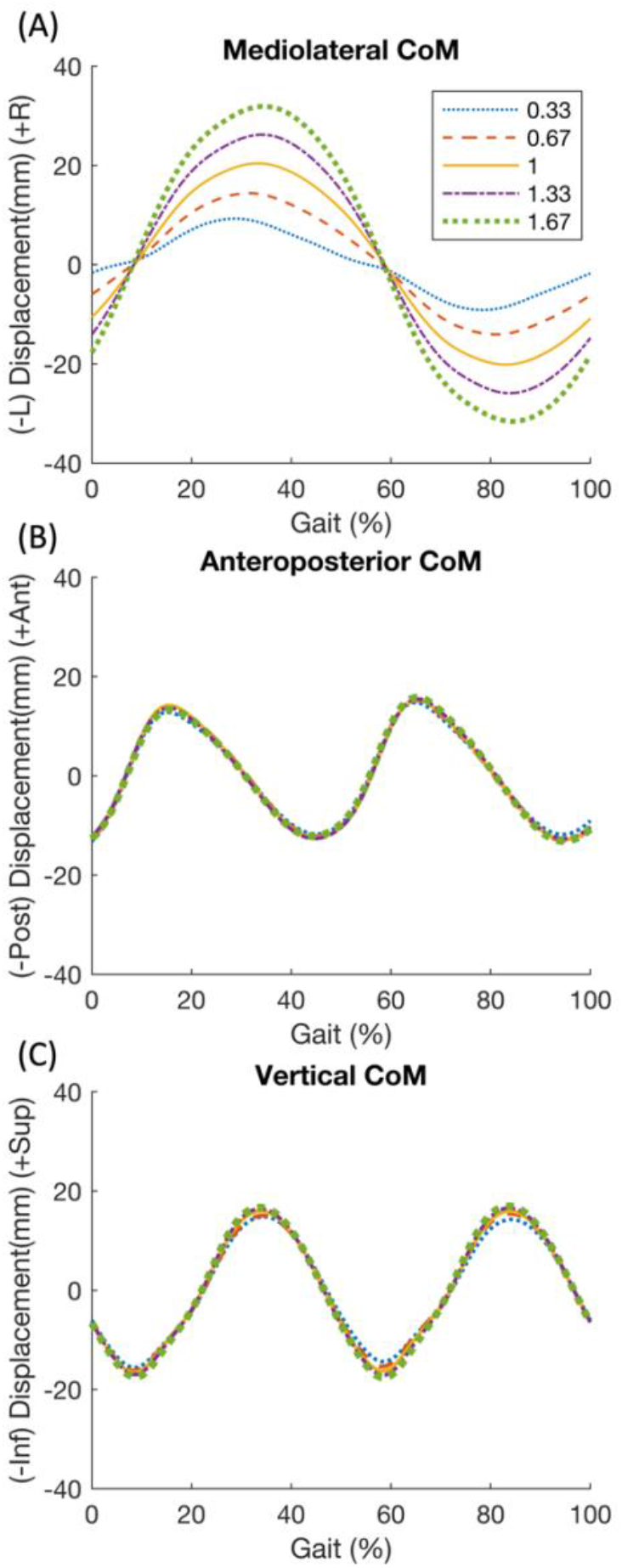
Group mean center of mass (CoM) in 5 prescribed step widths over gait cycle in the (A) mediolateral direction (L: left, R: right) (B) anteroposterior direction (Ant: anterior, Post: posterior), and (C) vertical direction (Sup: superior, Inf: inferior).

**Figure 5.**
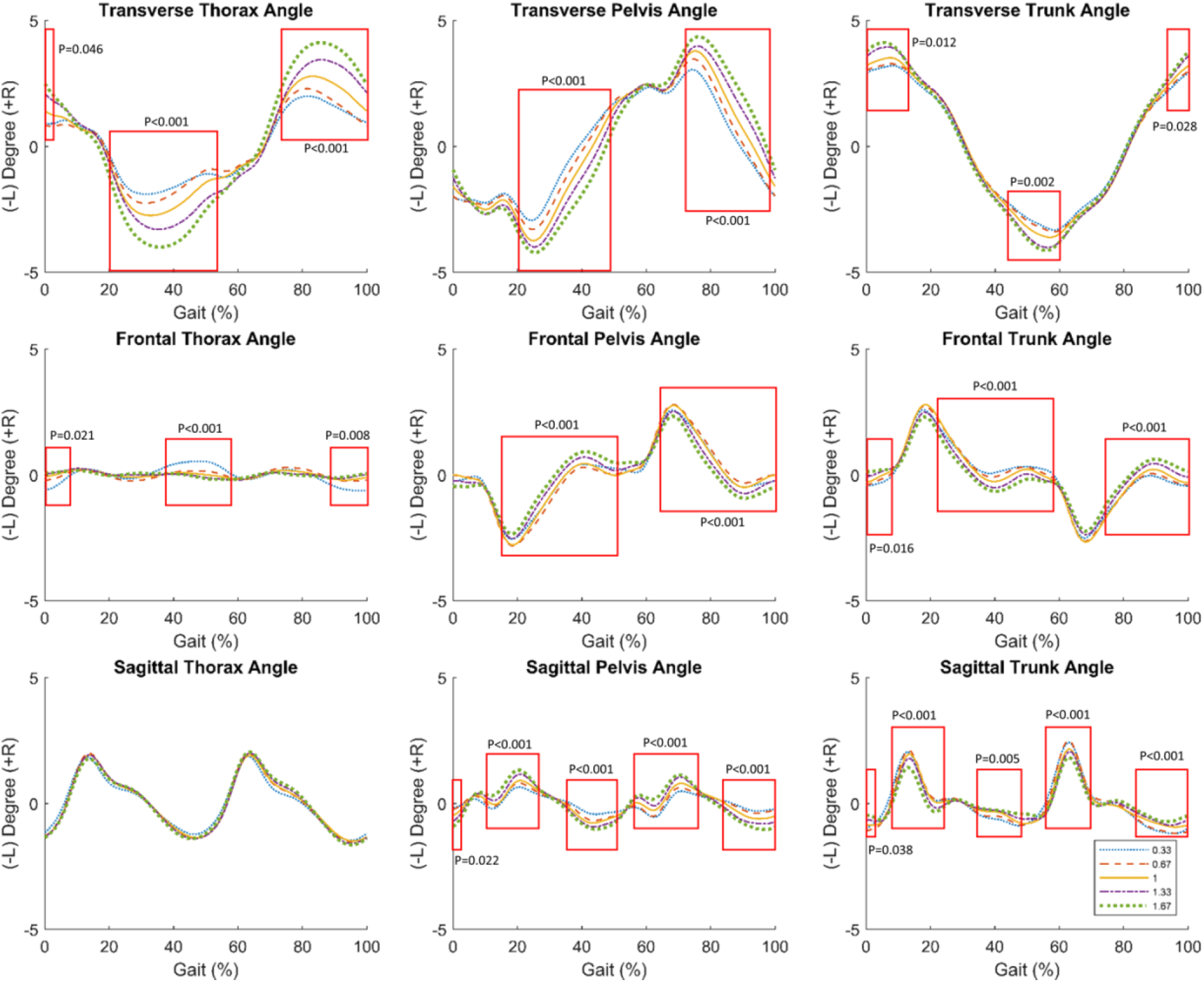
Group mean thorax, pelvis, and trunk angles in 5 prescribed step widths over gait cycle in the transverse, frontal and sagittal planes. Boxes highlight regions of significant oneway repeated measures ANOVA revealed by SPM analyses that was more than 10% gait cycle and consistently occurred during both left and right steps.

### Thorax, Pelvis, and Trunk Angular Excursions

Changes in step width affected pelvis, thorax, and trunk angular excursions most notably in the transverse plane, less so in the sagittal plane, and very little in the frontal plane (Fig 6). Statistical differences between different step widths for angular excursions were seen in the transverse plane thorax, pelvis, and trunk, and the sagittal plane pelvis and trunk (supplementary table).

**Figure 6.**
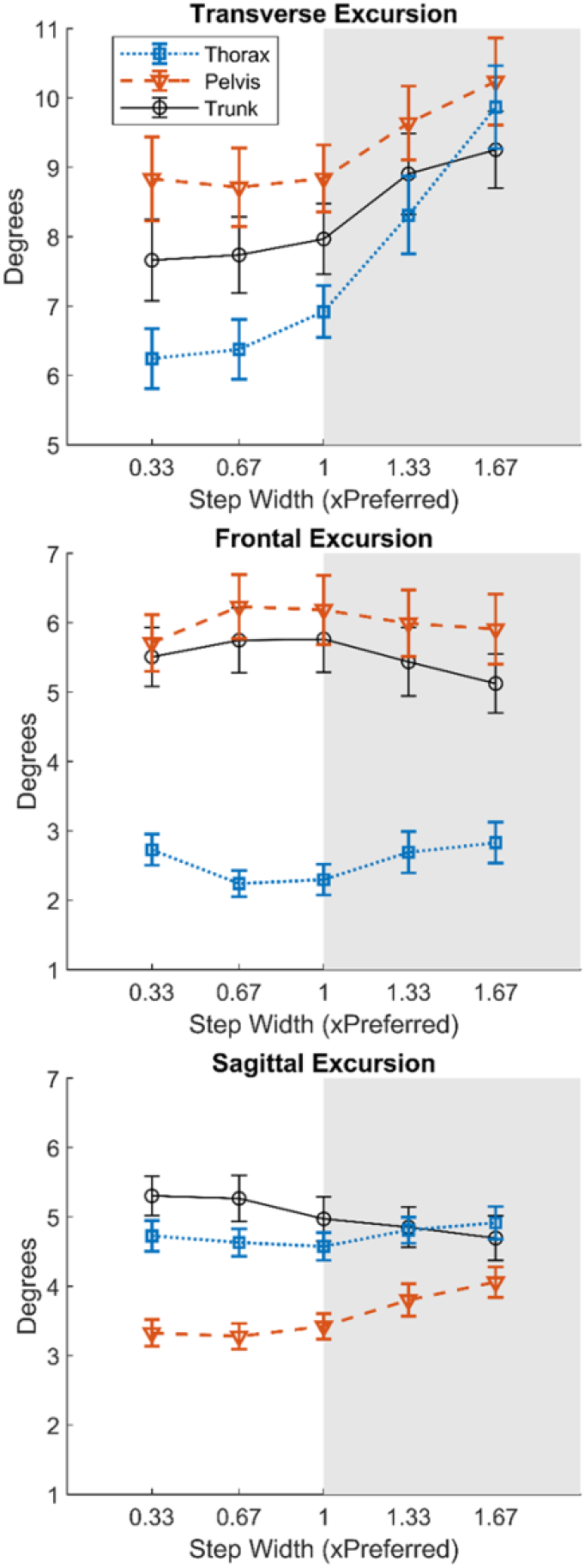
Mean and standard deviation of angular excursion of the thorax, pelvis, and trunk in the transverse, frontal, and sagittal planes for 5 prescribed step widths. Shaded background denotes step widths wider than preferred.

### Thorax-pelvis kinematic coordination

To characterize changes in thorax-pelvis coordination, we measured the cumulative proportion of time spent in four different patterns during the gait cycle: in-phase, anti-phase, pelvis-only, and thorax-only (Fig 7). In the transverse plane, thorax-pelvis coordination pattern consisted primarily of in-phase and pelvis-only pattern, where the segments either rotate towards the same direction or the pelvis rotate independently while the thorax remain stable. In the frontal plane, coordination was dominated by the pelvis-only pattern. In the sagittal plane, there was no clear dominant pattern.

**Figure 7.**
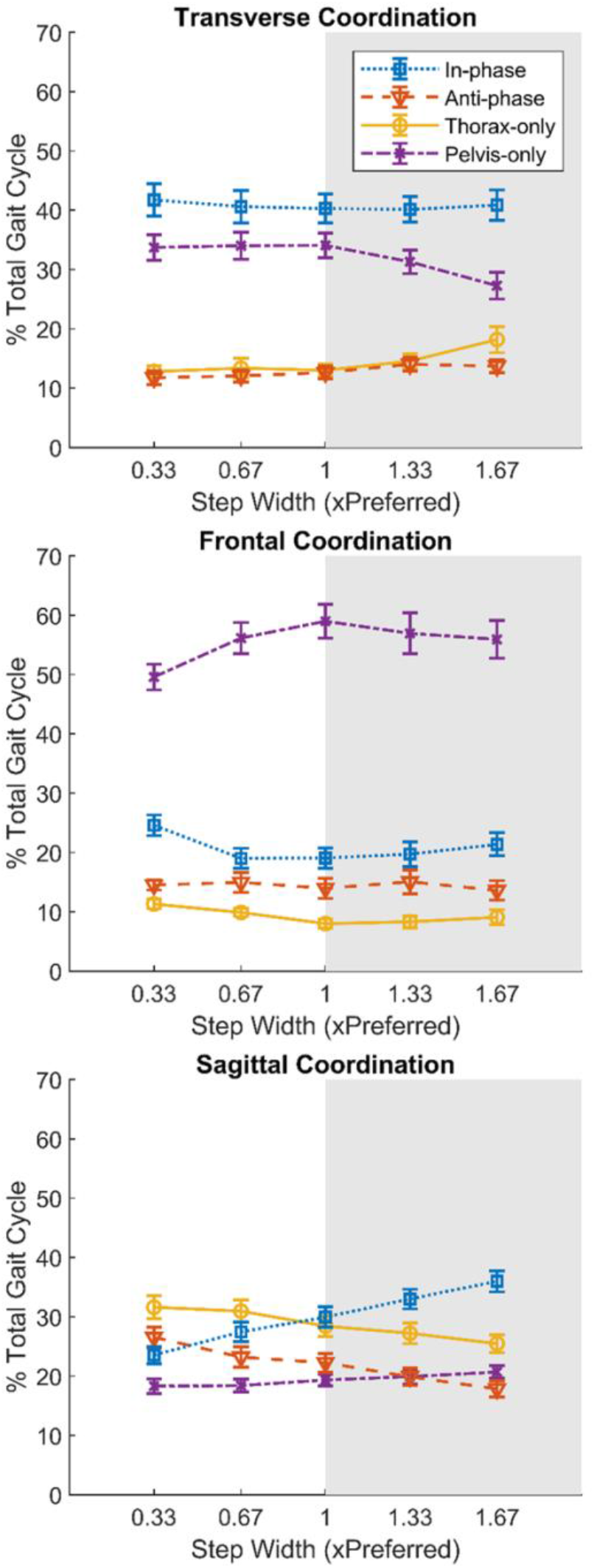
Mean and standard deviation of thorax-pelvis kinematic coordination in the transverse, frontal, and sagittal planes for 5 prescribed step widths. Shaded background denotes step widths wider than preferred.

Changes in thorax-pelvis kinematic coordination as a function of step width were more apparent at wider step widths in the transverse plane and narrower step widths in the frontal plane (Fig 7). Coordination was similarly affected by wider and narrower widths in the sagittal plane. Please refer to the supplementary table for statistical results.

### Longissimus Activation and Co-activation

Both peak longissimus activation and bilateral longissimus co-activation increased at narrower step widths, while at wider widths, bilateral co-activation decreased more markedly than peak activation (Fig 8). The peak right longissimus activation during 0.33PSW was significantly greater than 0.67, 1, 1.33 and 1.67PSW (p<0.025). The peak left longissimus activation during 0.33PSW was greater than 1PSW, 1.33PSW, and 1.67PSW (p<0.001), and activation at 0.67PSW was greater than 1.67PSW (p=0.003). Bilateral longissimus co-activation at 1.67PSW was less than 0.33, 0.67, and 1PSW (p<0.007), and co-activation at 1.33PSW was also less than 0.33PSW (p=0.005).

**Figure 8.**
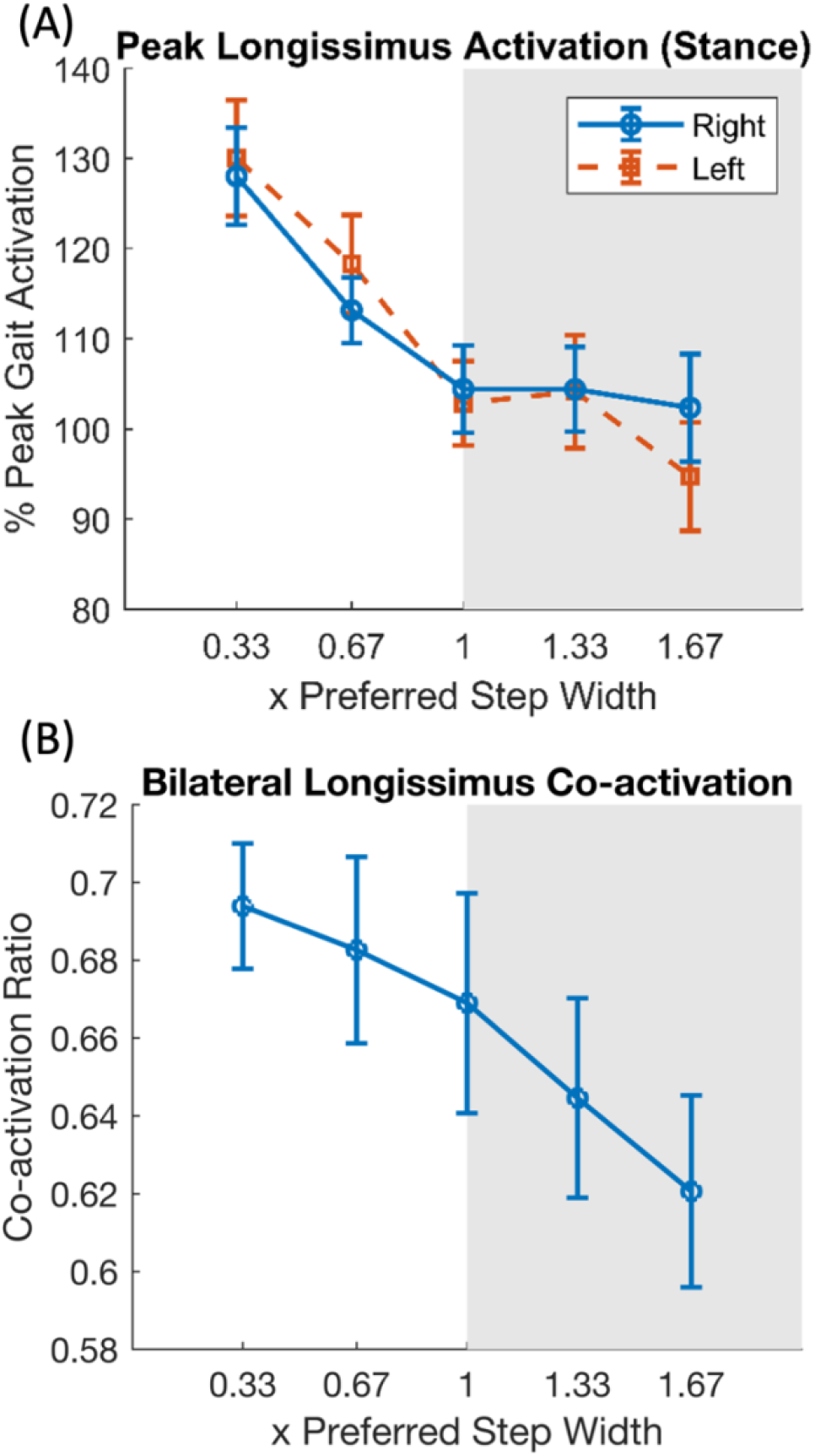
(A) Mean and standard deviation of the right and left longissimus peak activation during the contralateral stance phase. (B) Mean and standard deviation of bilateral co-activation ratio of the longissimus throughout the whole gait cycle.

## Discussion

We investigated the effects of varying step width on trunk control and specifically compared the effects of wider and narrower step widths. We found that although CoM only varied along the mediolateral axis across step widths, trunk kinematics in all three planes were affected. The results indicated that walking with wide and narrow step widths may be viewed as two distinct tasks, where greater stability demands were present in narrow widths and greater mechanical demands were present in wider widths. Angular excursions of the thorax, pelvis, and trunk were more affected by wider widths than narrower widths. Thorax-pelvis kinematic coordination was more affected by wider widths in transverse plane and by narrower widths in the frontal plane. Also, peak longissimus activation was more affected by narrower step widths, resulting in an increased activation. Bilateral longissimus co-activation decreased as step widths became wider.

In this study, participants’ average preferred step width was wider than the previously reported value of 0.13 x leg length (Donelan et al., 2001), probably due to the different methods of step width calculation (foot position in this study, center of pressure in Donelan et al, 2001). Participants closely adhered to the prescribed step widths, and the deviations from the prescribed step width at narrower widths were small and likely due to increased postural challenges and the range effect (a phenomenon in which the target at the extreme ranges is under-reached (Poulton, 1975)). Constant error increased as step width narrowed, but remained small nonetheless (<20% PSW). Variable error was small and consistent across step width. The participants’ consistent accuracy in matching of step widths to targets allowed us to analyze trunk control variables as a function of prescribed step width.

The systematic changes in the CoM displacement as a function of step width along the mediolateral axis reflected the constraints of the task. It is noteworthy, however, that CoM was invariant along the anterior-posterior and vertical axes, even though the trunk segments varied in all three planes. Thus, the motor system achieves a tight regulation of the CoM trajectory by multi-planar adjustments of the thorax and pelvis segments. The relationship between foot placement and CoM has been documented in previous studies (Arvin et al., 2016a; McAndrew Young and Dingwell, 2012; Perry and Srinivasan, 2017). Other studies have found that the mechanical state of the CoM during the swing phase predicts subsequent foot placement (Bruijn and van Dieën, 2018; Hurt et al., 2010; Stimpson et al., 2018; Wang and Srinivasan, 2014). However, none of these studies reported CoM behavior along axes other than mediolateral, nor did they describe detailed trunk kinematics. Our findings showed that thorax, pelvis, and trunk angles were adjusted across multiple planes, potentially due to coupling motion of the spinal structure (Barnes et al., 2009; Legaspi and Edmond, 2007). Furthermore, the multi-planar adjustments of trunk kinematics could also be an exploitation of motor redundancy, where different combinations of body segment configurations can produce the same CoM trajectory (Bernstein, 1967; Fietzer et al., 2019; Latash et al., 2010).

The variation in angular excursion of the thorax, pelvis, and trunk at different step widths was most apparent in the transverse plane. This is notable because step width was manipulated in the frontal plane. The time-series data (Fig 5) showed that although differences in kinematics as a function of step width did occur in the frontal plane, they occurred predominantly during mid-ranges of movement, and thus did not influence overall excursion. Additionally, transverse plane rotation is the dominant motion of the trunk during walking.

Transverse plane trunk rotation is associated with step length and arm swings (Pontzer et al., 2009; Saunders et al., 1953). We observed that when walking with wider step widths, participants demonstrated larger arm swings and greater diagonal step distance to maintain step length and keep up with the fixed treadmill speed. At narrower step widths, diagonal step distance does not become proportionally smaller based on geometric principles, which may have prevented trunk transverse plane excursions from further decreasing.

The thorax-pelvis coordination patterns changed with step width, more so during wide widths in the transverse plane and more so during the narrow widths in the frontal plane. Consistent with previous literature (Seay et al., 2011), in-phase and pelvis-only patterns dominated the transverse plane. The pelvis-only pattern dominated the frontal plane during walking since the thorax needed to remain level to stabilize the head for visual input. This is the first study to investigate how trunk coordination patterns change as step widths changes. In the transverse plane, when step width was wider than preferred, the proportion of the pelvis-only pattern decreased, and the proportion of the thorax-only pattern increased. In the frontal plane, the proportion of the pelvis-only pattern decreased, and the proportion of the in-phase pattern increased with narrow widths.

The changes in frontal plane coordination at narrow widths corresponded to an increased bilateral longissimus co-activation and demonstrated that increased active control is required with a smaller base of support (MacKinnon and Winter, 1993; Perry and Srinivasan, 2017). Peak longissimus activation and bilateral longissimus co-activation both increased as step width decreased. The paraspinal muscles are known to be controlled through a feedforward mechanism in a top-down manner from cervical to lumbar before heel strike during walking (Prince et al., 1994). The paraspinal muscles at the lumbar region have been shown to be the most highly activated (Prince et al., 1994), likely due to their critical location and the need to control the large mass and moment arm of the upper body. Donelan and colleagues showed evidence of metabolic cost necessary for active stabilization at narrow widths (Donelan et al., 2004). Our findings of increased muscle activation might be one of the contributors to the metabolic cost.

Although peak longissimus activation increased as step width narrowed from preferred width, peak activation was little affected by widening of the step width. One possible explanation is that the peak muscle activation not only contributes to increased active lateral stabilization at narrow widths but also counteracts the flexion moment of the trunk consistently present during heel strike at all widths. On the other hand, bilateral longissimus co-activation followed a linear trend and continued to decrease from preferred to wider widths, perhaps because it was a frontal plane-specific measurement that primarily captured lateral stabilization.

The current study had several limitations. Participants were a convenient sample of students who are young and active, which may limit generalizability to other populations. It is possible that participant did not reach a plateau of their preferred step width during the 3-minute familiarization trial. Acclimatization for step width during treadmill walking has been reported to be around 4.5 minutes, although most changes occur during the first 100 seconds (Meyer et al., 2019). Some statistically significant changes in trunk angular excursions or thorax-pelvis kinematic coordination were arguably small across step widths (for example, less than 2 degrees in pelvis transverse excursion). Nevertheless, the changes were relatively large when considering the total excursion or per cent coordination pattern used in walking, and the measurement had good test-retest reliability. This research provides a foundation for future studies on trunk control in various step widths on different populations, such as individuals with low back pain.

In summary, step width affected trunk control in various ways. Trunk kinematics and thorax-pelvis coordination in all three planes were affected by step width, with transverse plane variables greater impacted by wider widths and frontal plane variables greater impacted by narrower widths. Longissimus activation and bilateral co-activation both increased with narrow step widths, and peak activation showed greater changes with narrow step widths. Findings of this study support the hypothesis that wider and narrow step widths each present unique demand on trunk control. The results afford insight into how the trunk and pelvis segments are controlled to maintain stability at various step widths.

## Conflict of interest statement

The authors disclose no conflict of interest that could inappropriately influence this work.

## Acknowledgements

This research was supported by the International Society of Biomechanics Matching Dissertation Grant. The funding agency does not have any involvement in the design, execution, and the interpretation and write up of this study.

## Notes

### Competing Interest Statement

The authors have declared no competing interest.

### Summary of Updates

Minor revisions focusing on limitations of potentially inadequate familiarization period for the acclimation of step width on treadmill. Slight wording change, and one additional reference.

