## supplementary table for "Trunk Control during Gait: Walking with Wide and Narrow Step Widths Present Distinct Challenges"

**Supplementary Table.** Pairwise comparisons between step widths for all variables.

| **Variable** | **Step Width** | **SW=0.33** | **SW=0.67** | **SW=1** | **SW=1.33** | **SW=1.67** |
| --- | --- | --- | --- | --- | --- | --- |
| **Constant SW Error** | 0.33 | -- | **0.003** | **<0.001** | **<0.001** | **<0.001** |
|  | 0.67 |  | -- | 0.347 | 0.062 | **<0.001** |
|  | 1 |  |  | -- | 0.931 | 0.060 |
|  | 1.33 |  |  |  | -- | 0.338 |
|  | 1.67 |  |  |  |  | -- |
| **Variable SW Error** | 0.33 | -- | **0.014** | **0.001** | **0.004** | 0.073 |
|  | 0.67 |  | -- | 0.972 | 0.996 | 0.980 |
|  | 1 |  |  | -- | 0.999 | 0.760 |
|  | 1.33 |  |  |  | -- | 0.880 |
|  | 1.67 |  |  |  |  | -- |
| **Transverse Thorax Excursion** | 0.33 | -- | 0.998 | 0.455 | **<0.001** | **<0.001** |
|  | 0.67 |  | -- | 0.667 | **<0.001** | **<0.001** |
|  | 1 |  |  | -- | **0.006** | **<0.001** |
|  | 1.33 |  |  |  | -- | **0.001** |
|  | 1.67 |  |  |  |  | -- |
| **Transverse Pelvis Excursion** | 0.33 | -- | 0.999 | 1.000 | 0.390 | **0.018** |
|  | 0.67 |  | -- | 0.999 | 0.248 | **0.007** |
|  | 1 |  |  | -- | 0.401 | **0.018** |
|  | 1.33 |  |  |  | -- | 0.683 |
|  | 1.67 |  |  |  |  | -- |
| **Transverse Trunk Excursion** | 0.33 | -- | 0.999 | 0.850 | **<0.001** | **<0.001** |
|  | 0.67 |  | -- | 0.943 | **0.001** | **<0.001** |
|  | 1 |  |  | -- | **0.017** | **<0.001** |
|  | 1.33 |  |  |  | -- | 0.781 |
|  | 1.67 |  |  |  |  | -- |
| **Frontal Thorax Excursion^#^** | 0.33 | -- | na | na | na | na |
|  | 0.67 |  | -- | na | na | na |
|  | 1 |  |  | -- | na | na |
|  | 1.33 |  |  |  | -- | na |
|  | 1.67 |  |  |  |  | -- |
| **Frontal Pelvis Excursion^#^** | 0.33 | -- | na | na | na | na |
|  | 0.67 |  | -- | na | na | na |
|  | 1 |  |  | -- | na | na |
|  | 1.33 |  |  |  | -- | na |
|  | 1.67 |  |  |  |  | -- |
| **Frontal Trunk Excursion^#^** | 0.33 | -- | na | na | na | na |
|  | 0.67 |  | -- | na | na | na |
|  | 1 |  |  | -- | na | na |
|  | 1.33 |  |  |  | -- | na |
|  | 1.67 |  |  |  |  | -- |
| **Sagittal Thorax Excursion^#^** | 0.33 | -- | na | na | na | na |
|  | 0.67 |  | -- | na | na | na |
|  | 1 |  |  | -- | na | na |
|  | 1.33 |  |  |  | -- | na |
|  | 1.67 |  |  |  |  | -- |
| **Sagittal**  **Pelvis**  **Excursion** | 0.33 | -- | 0.996 | 0.953 | **0.003** | **<0.001** |
|  | 0.67 |  | -- | 0.815 | **<0.001** | **<0.001** |
|  | 1 |  |  | -- | **0.035** | **<0.001** |
|  | 1.33 |  |  |  | -- | 0.297 |
|  | 1.67 |  |  |  |  | -- |
| **Sagittal Trunk Excursion** | 0.33 | -- | 0.999 | 0.222 | **0.036** | **0.001** |
|  | 0.67 |  | -- | 0.335 | 0.067 | **0.003** |
|  | 1 |  |  | -- | 0.944 | 0.404 |
|  | 1.33 |  |  |  | -- | 0.857 |
|  | 1.67 |  |  |  |  | -- |
| **Transverse In-phase^#^** | 0.33 | -- | na | na | na | na |
|  | 0.67 |  | -- | na | na | na |
|  | 1 |  |  | -- | na | na |
|  | 1.33 |  |  |  | -- | na |
|  | 1.67 |  |  |  |  | -- |
| **Transverse Anti-phase** | 0.33 | -- | 0.995 | 0.808 | **0.041** | 0.111 |
|  | 0.67 |  | -- | 0.957 | 0.115 | 0.258 |
|  | 1 |  |  | -- | 0.433 | 0.680 |
|  | 1.33 |  |  |  | -- | 0.995 |
|  | 1.67 |  |  |  |  | -- |
| **Transverse Thorax-only** | 0.33 | -- | 0.997 | 1.000 | 0.810 | **0.007** |
|  | 0.67 |  | -- | 1.000 | 0.944 | **0.023** |
|  | 1 |  |  | -- | 0.874 | **0.012** |
|  | 1.33 |  |  |  | -- | 0.163 |
|  | 1.67 |  |  |  |  | -- |
| **Transverse Pelvis-only** | 0.33 | -- | 1.000 | 1.000 | 0.685 | **0.005** |
|  | 0.67 |  | -- | 1.000 | 0.589 | **0.003** |
|  | 1 |  |  | -- | 0.565 | **0.002** |
|  | 1.33 |  |  |  | -- | 0.191 |
|  | 1.67 |  |  |  |  | -- |
| **Frontal In-phase** | 0.33 | -- | **0.002** | **0.002** | **0.009** | 0.197 |
|  | 0.67 |  | -- | 1.000 | 0.990 | 0.498 |
|  | 1 |  |  | -- | 0.992 | 0.517 |
|  | 1.33 |  |  |  | -- | 0.794 |
|  | 1.67 |  |  |  |  | -- |
| **Frontal Anti-phase^#^** | 0.33 | -- | na | na | na | na |
|  | 0.67 |  | -- | na | na | na |
|  | 1 |  |  | -- | na | na |
|  | 1.33 |  |  |  | -- | na |
|  | 1.67 |  |  |  |  | -- |
| **Frontal Thorax-only** | 0.33 | -- | 0.655 | **0.015** | **0.036** | 0.217 |
|  | 0.67 |  | -- | 0.393 | 0.570 | 0.944 |
|  | 1 |  |  | -- | 0.999 | 0.848 |
|  | 1.33 |  |  |  | -- | 0.947 |
|  | 1.67 |  |  |  |  | -- |
| **Frontal Pelvis-only** | 0.33 | -- | **0.033** | **<0.001** | **0.011** | **0.044** |
|  | 0.67 |  | -- | 0.730 | 0.997 | 1.000 |
|  | 1 |  |  | -- | 0.899 | 0.669 |
|  | 1.33 |  |  |  | -- | 0.992 |
|  | 1.67 |  |  |  |  | -- |
| **Sagittal In-phase** | 0.33 | -- | **0.023** | **<0.001** | **<0.001** | **<0.001** |
|  | 0.67 |  | -- | 0.304 | **<0.001** | **<0.001** |
|  | 1 |  |  | -- | 0.136 | **<0.001** |
|  | 1.33 |  |  |  | -- | 0.143 |
|  | 1.67 |  |  |  |  | -- |
| **Sagittal Anti-phase** | 0.33 | -- | **0.007** | **<0.001** | **<0.001** | **<0.001** |
|  | 0.67 |  | -- | 0.876 | **0.007** | **<0.001** |
|  | 1 |  |  | -- | 0.116 | **<0.001** |
|  | 1.33 |  |  |  | -- | 0.230 |
|  | 1.67 |  |  |  |  | -- |
| **Sagittal Thorax-only** | 0.33 | -- | 0.980 | 0.055 | **0.002** | **<0.001** |
|  | 0.67 |  | -- | 0.209 | **0.014** | **<0.001** |
|  | 1 |  |  | -- | 0.844 | 0.092 |
|  | 1.33 |  |  |  | -- | 0.582 |
|  | 1.67 |  |  |  |  | -- |
| **Sagittal Pelvis-only** | 0.33 | -- | 1.000 | 0.763 | 0.376 | 0.056 |
|  | 0.67 |  | -- | 0.821 | 0.441 | 0.075 |
|  | 1 |  |  | -- | 0.973 | 0.559 |
|  | 1.33 |  |  |  | -- | 0.901 |
|  | 1.67 |  |  |  |  | -- |
| **Right Longissimus Peak Activation** | 0.33 | -- | **0.025** | **<0.001** | **<0.001** | **<0.001** |
|  | 0.67 |  | -- | 0.408 | 0.406 | 0.197 |
|  | 1 |  |  | -- | 1.000 | 0.994 |
|  | 1.33 |  |  |  | -- | 0.994 |
|  | 1.67 |  |  |  |  | -- |
| **Left Longissimus Peak Activation** | 0.33 | -- | 0.370 | **<0.001** | **<0.001** | **<0.001** |
|  | 0.67 |  | -- | 0.122 | 0.190 | **0.003** |
|  | 1 |  |  | -- | 1.000 | 0.724 |
|  | 1.33 |  |  |  | -- | 0.597 |
|  | 1.67 |  |  |  |  | -- |
| **Longissimus Co-contraction** | 0.33 | -- | 0.935 | 0.413 | **0.005** | **<0.001** |
|  | 0.67 |  | -- | 0.878 | 0.063 | **<0.001** |
|  | 1 |  |  | -- | 0.436 | **0.007** |
|  | 1.33 |  |  |  | -- | 0.453 |
|  | 1.67 |  |  |  |  | -- |

^#^One-way repeated measures ANOVA F-test not significant.

**Table 2.** Constant step width error pairwise comparison (ANOVA F=12.96, p<0.001).

| **Step Width (I)** | **Step Width (J)** | **Mean difference (J-I)** | **Std. Error** | **P-value** |
| --- | --- | --- | --- | --- |
| **0.33** | 0.67 | -9.970^*^ | 2.483 | <0.001 |
|  | 1 | -14.867^*^ | 2.483 | <0.001 |
|  | 1.33 | -14.059^*^ | 2.483 | <0.001 |
|  | 1.67 | -14.705^*^ | 2.483 | <0.001 |
| **0.67** | 1 | -4.898 | 2.483 | 0.279 |
|  | 1.33 | -4.090 | 2.483 | 0.467 |
|  | 1.67 | -4.736 | 2.483 | 0.313 |
| **1** | 1.33 | 0.808 | 2.483 | 0.998 |
|  | 1.67 | 0.162 | 2.483 | 1.000 |
| **1.33** | 1.67 | -0.646 | 2.483 | 0.999 |

^*^Significant at the 0.05 level

**Table 3.** Variable step width error pairwise comparison (ANOVA F=4.65, p=0.0021).

| **Step Width (I)** | **Step Width (J)** | **Mean difference (J-I)** | **Std. Error** | **P-value** |
| --- | --- | --- | --- | --- |
| **0.33** | 0.67 | -3.850^*^ | 1.222 | 0.014 |
|  | 1 | -4.609^*^ | 1.222 | 0.002 |
|  | 1.33 | -4.313^*^ | 1.222 | 0.003 |
|  | 1.67 | -3.161 | 1.222 | 0.073 |
| **0.67** | 1 | -0.758 | 1.222 | 0.972 |
|  | 1.33 | -0.463 | 1.222 | 0.996 |
|  | 1.67 | 0.689 | 1.222 | 0.980 |
| **1** | 1.33 | 0.296 | 1.222 | 0.999 |
|  | 1.67 | 1.447 | 1.222 | 0.760 |
| **1.33** | 1.67 | 1.15 | 1.222 | 0.880 |

^*^Significant at the 0.05 level

**Table 4.** Transverse plane thorax excursion pairwise comparison (ANOVA F=28.50, p<0.0001).

| **Step Width (I)** | **Step Width (J)** | **Mean difference (J-I)** | **Std. Error** | **P-value** |
| --- | --- | --- | --- | --- |
| **0.33** | 0.67 | 0.133 | 0.407 | 0.998 |
|  | 1 | 0.678 | 0.407 | 0.455 |
|  | 1.33 | 2.068^*^ | 0.407 | <0.001 |
|  | 1.67 | 3.627^*^ | 0.407 | <0.001 |
| **0.67** | 1 | 0.545 | 0.407 | 0.667 |
|  | 1.33 | 1.935^*^ | 0.407 | <0.001 |
|  | 1.67 | 3.494^*^ | 0.407 | <0.001 |
| **1** | 1.33 | 1.390^*^ | 0.407 | 0.006 |
|  | 1.67 | 2.949^*^ | 0.407 | <0.001 |
| **1.33** | 1.67 | 1.559^*^ | 0.407 | 0.001 |

^*^Significant at the 0.05 level

**Table 5.** Transverse plane pelvis excursion pairwise comparison (ANOVA F=4.238, p=0.004).

| **Step Width (I)** | **Step Width (J)** | **Mean difference (J-I)** | **Std. Error** | **P-value** |
| --- | --- | --- | --- | --- |
| **0.33** | 0.67 | -0.121 | 0.456 | 0.999 |
|  | 1 | 0.008 | 0.456 | 1.000 |
|  | 1.33 | 0.808 | 0.456 | 0.390 |
|  | 1.67 | 1.406^*^ | 0.456 | 0.017 |
| **0.67** | 1 | 0.129 | 0.456 | 0.999 |
|  | 1.33 | 0.929 | 0.456 | 0.248 |
|  | 1.67 | 1.527^*^ | 0.456 | 0.007 |
| **1** | 1.33 | 0.799 | 0.456 | 0.401 |
|  | 1.67 | 1.398^*^ | 0.456 | 0.018 |
| **1.33** | 1.67 | 0.599 | 0.456 | 0.683 |

^*^Significant at the 0.05 level

**Table 6.** Transverse plane trunk excursion pairwise comparison (ANOVA F=11.50, p=<0.0001).

| **Step Width (I)** | **Step Width (J)** | **Mean difference (J-I)** | **Std. Error** | **P-value** |
| --- | --- | --- | --- | --- |
| **0.33** | 0.67 | 0.077 | 0.303 | 0.999 |
|  | 1 | 0.307 | 0.303 | 0.850 |
|  | 1.33 | 1.246^*^ | 0.303 | <0.001 |
|  | 1.67 | 1.594^*^ | 0.303 | <0.001 |
| **0.67** | 1 | 0.230 | 0.303 | 0.943 |
|  | 1.33 | 1.169^*^ | 0.303 | 0.001 |
|  | 1.67 | 1.517^*^ | 0.303 | <0.001 |
| **1** | 1.33 | 0.940^*^ | 0.303 | 0.017 |
|  | 1.67 | 1.287^*^ | 0.303 | <0.001 |
| **1.33** | 1.67 | 0.348^*^ | 0.303 | 0.781 |

^*^Significant at the 0.05 level

**Table 7.** Sagittal plane pelvis excursion pairwise comparison (ANOVA F=12.977, p<0.001).

| **Step Width (I)** | **Step Width (J)** | **Mean difference (J-I)** | **Std. Error** | **P-value** |
| --- | --- | --- | --- | --- |
| **0.33** | 0.67 | -0.049 | 0.133 | 0.996 |
|  | 1 | 0.095 | 0.133 | 0.953 |
|  | 1.33 | 0.475^*^ | 0.133 | 0.003 |
|  | 1.67 | 0.732^*^ | 0.133 | <0.001 |
| **0.67** | 1 | 0.144 | 0.133 | 0.815 |
|  | 1.33 | 0.524^*^ | 0.133 | <0.001 |
|  | 1.67 | 0.781^*^ | 0.133 | <0.001 |
| **1** | 1.33 | 0.380^*^ | 0.133 | 0.035 |
|  | 1.67 | 0.637^*^ | 0.133 | <0.001 |
| **1.33** | 1.67 | 0.258 | 0.133 | 0.297 |

^*^Significant at the 0.05 level

**Table 8.** Sagittal plane trunk excursion pairwise comparison (ANOVA F=5.534, p<0.001).

| **Step Width (I)** | **Step Width (J)** | **Mean difference (J-I)** | **Std. Error** | **P-value** |
| --- | --- | --- | --- | --- |
| **0.33** | 0.67 | -0.036 | 0.158 | 0.999 |
|  | 1 | -0.331 | 0.158 | 0.222 |
|  | 1.33 | -0.451^*^ | 0.158 | 0.036 |
|  | 1.67 | -0.609^*^ | 0.158 | 0.001 |
| **0.67** | 1 | -0.296 | 0.158 | 0.335 |
|  | 1.33 | -0.415^*^ | 0.158 | 0.067 |
|  | 1.67 | -0.572^*^ | 0.158 | 0.003 |
| **1** | 1.33 | -0.119 | 0.158 | 0.944 |
|  | 1.67 | -0.277 | 0.158 | 0.404 |
| **1.33** | 1.67 | -0.158 | 0.158 | 0.857 |

^*^Significant at the 0.05 level

**Table 9.** Transverse plane anti-phase coordination pairwise comparison (ANOVA F=3.014, p=0.023).

| **Step Width (I)** | **Step Width (J)** | **Mean difference (J-I)** | **Std. Error** | **P-value** |
| --- | --- | --- | --- | --- |
| **0.33** | 0.67 | 0.323 | 0.808 | 0.995 |
|  | 1 | 0.888 | 0.808 | 0.808 |
|  | 1.33 | 2.263^*^ | 0.808 | 0.041 |
|  | 1.67 | 1.953 | 0.808 | 0.111 |
| **0.67** | 1 | 0.565 | 0.808 | 0.957 |
|  | 1.33 | 1.940 | 0.808 | 0.115 |
|  | 1.67 | 1.630 | 0.808 | 0.258 |
| **1** | 1.33 | 1.375 | 0.808 | 0.433 |
|  | 1.67 | 1.065 | 0.808 | 0.680 |
| **1.33** | 1.67 | -0.310 | 0.808 | 0.995 |

^*^Significant at the 0.05 level

**Table 10.** Transverse plane thorax-only coordination pairwise comparison (ANOVA F=3.830, p=0.007).

| **Step Width (I)** | **Step Width (J)** | **Mean difference (J-I)** | **Std. Error** | **P-value** |
| --- | --- | --- | --- | --- |
| **0.33** | 0.67 | 0.553 | 1.615 | 0.997 |
|  | 1 | 0.220 | 1.615 | 1.000 |
|  | 1.33 | 1.765 | 1.615 | 0.810 |
|  | 1.67 | 5.393^*^ | 1.615 | 0.008 |
| **0.67** | 1 | -0.333 | 1.615 | 1.000 |
|  | 1.33 | 1.213 | 1.615 | 0.944 |
|  | 1.67 | 4.840^*^ | 1.615 | 0.023 |
| **1** | 1.33 | 1.545 | 1.615 | 0.874 |
|  | 1.67 | 5.173^*^ | 1.615 | 0.012 |
| **1.33** | 1.67 | 3.628 | 1.615 | 0.163 |

^*^Significant at the 0.05 level

**Table 11.** Transverse plane pelvis-only coordination pairwise comparison (ANOVA F=4.955, p=0.001).

| **Step Width (I)** | **Step Width (J)** | **Mean difference (J-I)** | **Std. Error** | **P-value** |
| --- | --- | --- | --- | --- |
| **0.33** | 0.67 | 0.278 | 1.855 | 1.000 |
|  | 1 | 0.345 | 1.855 | 1.000 |
|  | 1.33 | -2.430 | 1.855 | 0.685 |
|  | 1.67 | -6.455^*^ | 1.855 | 0.005 |
| **0.67** | 1 | 0.068 | 1.855 | 1.000 |
|  | 1.33 | -2.708 | 1.855 | 0.589 |
|  | 1.67 | -6.733^*^ | 1.855 | 0.003 |
| **1** | 1.33 | -2.775 | 1.855 | 0.565 |
|  | 1.67 | -6.800^*^ | 1.855 | 0.002 |
| **1.33** | 1.67 | -4.025 | 1.855 | 0.191 |

^*^Significant at the 0.05 level

**Table 12.** Frontal plane in-phase coordination pairwise comparison (ANOVA F=5.007, p=0.001).

| **Step Width (I)** | **Step Width (J)** | **Mean difference (J-I)** | **Std. Error** | **P-value** |
| --- | --- | --- | --- | --- |
| **0.33** | 0.67 | -5.570^*^ | 1.484 | 0.002 |
|  | 1 | -5.528^*^ | 1.484 | 0.002 |
|  | 1.33 | -4.865^*^ | 1.484 | 0.009 |
|  | 1.67 | -3.198 | 1.484 | 0.197 |
| **0.67** | 1 | 0.043 | 1.484 | 1.000 |
|  | 1.33 | 0.705 | 1.484 | 0.990 |
|  | 1.67 | 2.373 | 1.484 | 0.498 |
| **1** | 1.33 | 0.663 | 1.484 | 0.992 |
|  | 1.67 | 2.330 | 1.484 | 0.517 |
| **1.33** | 1.67 | -1.668 | 1.484 | 0.794 |

^*^Significant at the 0.05 level

**Table 13.** Frontal plane thorax-only coordination pairwise comparison (ANOVA F=3.171, p=0.018).

| **Step Width (I)** | **Step Width (J)** | **Mean difference (J-I)** | **Std. Error** | **P-value** |
| --- | --- | --- | --- | --- |
| **0.33** | 0.67 | -1.438 | 1.059 | 0.655 |
|  | 1 | -3.310^*^ | 1.059 | 0.015 |
|  | 1.33 | -3.015^*^ | 1.059 | 0.036 |
|  | 1.67 | -2.233 | 1.059 | 0.217 |
| **0.67** | 1 | -1.873 | 1.059 | 0.393 |
|  | 1.33 | -1.578 | 1.059 | 0.570 |
|  | 1.67 | -0.795 | 1.059 | 0.944 |
| **1** | 1.33 | 0.295 | 1.059 | 0.999 |
|  | 1.67 | 1.078 | 1.059 | 0.848 |
| **1.33** | 1.67 | 0.783 | 1.059 | 0.947 |

^*^Significant at the 0.05 level

**Table 14.** Sagittal plane in-phase coordination pairwise comparison (ANOVA F=27.432, p<0.001).

| **Step Width (I)** | **Step Width (J)** | **Mean difference (J-I)** | **Std. Error** | **P-value** |
| --- | --- | --- | --- | --- |
| **0.33** | 0.67 | 3.902^*^ | 1.301 | 0.023 |
|  | 1 | 6.407^*^ | 1.301 | <0.001 |
|  | 1.33 | 9.435^*^ | 1.301 | <0.001 |
|  | 1.67 | 12.435^*^ | 1.301 | <0.001 |
| **0.67** | 1 | 2.505 | 1.301 | 0.304 |
|  | 1.33 | 5.533^*^ | 1.301 | <0.001 |
|  | 1.67 | 8.532^*^ | 1.301 | <0.001 |
| **1** | 1.33 | 3.028 | 1.301 | 0.136 |
|  | 1.67 | 6.027^*^ | 1.301 | <0.001 |
| **1.33** | 1.67 | 3.000 | 1.301 | 0.143 |

^*^Significant at the 0.05 level

**Table 15.** Sagittal plane anti-phase coordination pairwise comparison (ANOVA F=22.432, p<0.001).

| **Step Width (I)** | **Step Width (J)** | **Mean difference (J-I)** | **Std. Error** | **P-value** |
| --- | --- | --- | --- | --- |
| **0.33** | 0.67 | -3.330^*^ | 0.985 | 0.006 |
|  | 1 | -4.270^*^ | 0.985 | <0.001 |
|  | 1.33 | -6.633^*^ | 0.985 | <0.001 |
|  | 1.67 | -8.680^*^ | 0.985 | <0.001 |
| **0.67** | 1 | -0.940 | 0.985 | 0.876 |
|  | 1.33 | -3.303^*^ | 0.985 | 0.007 |
|  | 1.67 | -5.350^*^ | 0.985 | <0.001 |
| **1** | 1.33 | -2.363 | 0.985 | 0.116 |
|  | 1.67 | -4.410^*^ | 0.985 | <0.001 |
| **1.33** | 1.67 | -2.048 | 0.985 | 0.230 |

^*^Significant at the 0.05 level

**Table 16.** Sagittal plane thorax-only coordination pairwise comparison (ANOVA F=9.349, p<0.001).

| **Step Width (I)** | **Step Width (J)** | **Mean difference (J-I)** | **Std. Error** | **P-value** |
| --- | --- | --- | --- | --- |
| **0.33** | 0.67 | -0.668 | 1.182 | 0.980 |
|  | 1 | -3.180 | 1.182 | 0.055 |
|  | 1.33 | -4.128^*^ | 1.182 | 0.002 |
|  | 1.67 | -6.123^*^ | 1.182 | <0.001 |
| **0.67** | 1 | -2.513 | 1.182 | 0.209 |
|  | 1.33 | -3.723^*^ | 1.182 | 0.014 |
|  | 1.67 | -5.460^*^ | 1.182 | <0.001 |
| **1** | 1.33 | -1.210 | 1.182 | 0.844 |
|  | 1.67 | -2.948 | 1.182 | 0.092 |
| **1.33** | 1.67 | -1.738 | 1.182 | 0.582 |

^*^Significant at the 0.05 level

**Table 17.** Sagittal plane thorax-only coordination pairwise comparison (ANOVA F=2.591, p=0.043).

| **Step Width (I)** | **Step Width (J)** | **Mean difference (J-I)** | **Std. Error** | **P-value** |
| --- | --- | --- | --- | --- |
| **0.33** | 0.67 | 0.095 | 0.884 | 1.000 |
|  | 1 | 1.043 | 0.884 | 0.763 |
|  | 1.33 | 1.588 | 0.884 | 0.376 |
|  | 1.67 | 2.373 | 0.884 | 0.056 |
| **0.67** | 1 | 0.948 | 0.884 | 0.821 |
|  | 1.33 | 1.493 | 0.884 | 0.441 |
|  | 1.67 | 2.278 | 0.884 | 0.075 |
| **1** | 1.33 | 0.545 | 0.884 | 0.973 |
|  | 1.67 | 1.330 | 0.884 | 0.559 |
| **1.33** | 1.67 | 0.785 | 0.884 | 0.901 |

^*^Significant at the 0.05 level

**Table 18.** Right peak longissimus activation pairwise comparison (ANOVA F=9.049, p<0.001).

| **Step Width (I)** | **Step Width (J)** | **Mean difference (J-I)** | **Std. Error** | **P-value** |
| --- | --- | --- | --- | --- |
| **0.33** | 0.67 | -14.873^*^ | 5.012 | 0.025 |
|  | 1 | -23.601^*^ | 5.012 | <0.001 |
|  | 1.33 | -23.624^*^ | 5.012 | <0.001 |
|  | 1.67 | -25.673^*^ | 5.012 | <0.001 |
| **0.67** | 1 | -8.728 | 5.012 | 0.408 |
|  | 1.33 | -8.752 | 5.012 | 0.406 |
|  | 1.67 | -10.800 | 5.012 | 0.197 |
| **1** | 1.33 | -0.024 | 5.012 | 1.000 |
|  | 1.67 | -2.072 | 5.012 | 0.994 |
| **1.33** | 1.67 | -2.048 | 5.012 | 0.994 |

^*^Significant at the 0.05 level

**Table 19.** Left peak longissimus activation pairwise comparison (ANOVA F=9.318, p<0.001).

| **Step Width (I)** | **Step Width (J)** | **Mean difference (J-I)** | **Std. Error** | **P-value** |
| --- | --- | --- | --- | --- |
| **0.33** | 0.67 | -11.759 | 6.511 | 0.370 |
|  | 1 | -27.218^*^ | 6.511 | <0.001 |
|  | 1.33 | -25.909^*^ | 6.511 | <0.001 |
|  | 1.67 | -35.330^*^ | 6.511 | <0.001 |
| **0.67** | 1 | -15.458 | 6.511 | 0.122 |
|  | 1.33 | -14.149 | 6.511 | 0.190 |
|  | 1.67 | -23.570^*^ | 6.511 | 0.003 |
| **1** | 1.33 | 1.309 | 6.511 | 1.000 |
|  | 1.67 | -8.112 | 6.511 | 0.724 |
| **1.33** | 1.67 | -9.421 | 6.511 | 0.597 |

^*^Significant at the 0.05 level

**Table 20.** Bilateral longissimus co-activation pairwise comparison (ANOVA F=8.491, p<0.001).

| **Step Width (I)** | **Step Width (J)** | **Mean difference (J-I)** | **Std. Error** | **P-value** |
| --- | --- | --- | --- | --- |
| **0.33** | 0.67 | -0.011 | 0.014 | 0.945 |
|  | 1 | -0.025 | 0.014 | 0.413 |
|  | 1.33 | -0.049^*^ | 0.014 | 0.005 |
|  | 1.67 | -0.073^*^ | 0.014 | <0.001 |
| **0.67** | 1 | -0.014 | 0.014 | 0.878 |
|  | 1.33 | -0.038 | 0.014 | 0.063 |
|  | 1.67 | -0.062^*^ | 0.014 | <0.001 |
| **1** | 1.33 | -0.024 | 0.014 | 0.436 |
|  | 1.67 | -0.048^*^ | 0.014 | 0.007 |
| **1.33** | 1.67 | -0.024 | 0.014 | 0.453 |

^*^Significant at the 0.05 level
